# GenomeQC: A quality assessment tool for genome assemblies and gene structure annotations

**DOI:** 10.1101/795237

**Authors:** Nancy Manchanda, John L. Portwood, Margaret R. Woodhouse, Arun S. Seetharam, Carolyn J. Lawrence-Dill, Carson M. Andorf, Matthew B. Hufford

**Author notes:** To whom correspondence should be addressed. Carson M. Andorf, Matthew B. Hufford.

## Abstract

**Background:** Genome assemblies are foundational for understanding the biology of a species. They provide a physical framework for mapping additional sequences, thereby enabling characterization of, for example, genomic diversity and differences in gene expression across individuals and tissue types. Quality metrics for genome assemblies gauge both the completeness and contiguity of an assembly and help provide confidence in downstream biological insights. To compare quality across multiple assemblies, a set of common metrics are typically calculated and then compared to one or more gold standard reference genomes. While several tools exist for calculating individual metrics, applications providing comprehensive evaluations of multiple assembly features are, perhaps surprisingly, lacking. Here, we describe a new toolkit that integrates multiple metrics to characterize both assembly and gene annotation quality in a way that enables comparison across multiple assemblies and assembly types.

**Findings:** Our application, named GenomeQC, is an easy-to-use and interactive web framework that integrates various quantitative measures to characterize genome assemblies and annotations. GenomeQC provides researchers with a comprehensive summary of these statistics and allows for benchmarking against gold standard reference assemblies.

**Conclusions:** The GenomeQC web application is implemented in R/Shiny version 1.5.9 and Python 3.6 and is freely available at https://genomeqc.maizegdb.org/ under the GPL license.

All source code and a containerized version of the GenomeQC pipeline is available in the GitHub repository https://github.com/HuffordLab/GenomeQC.

## Background

Over the past few decades, numerous plant genome assemblies have been generated, ranging in size from 63 Mb in *Genlisea aurea* [1] to 22 Gb in *Pinus taeda* [2]. The genomic resources generated from such projects have contributed to the development of improved crop varieties, enhanced our understanding of genome size, architecture, and complexity, and uncovered mechanisms underlying plant growth and development [3][4]. With the declining cost of sequence, the number of genome assemblies has increased exponentially (Supplementary Figure 1). The NCBI assembly database [5] currently hosts more than 800 plant genome assemblies with varying degrees of contiguity and increasingly includes multiple genome assemblies per species (Supplementary Figure 2).

**Fig. 1.**
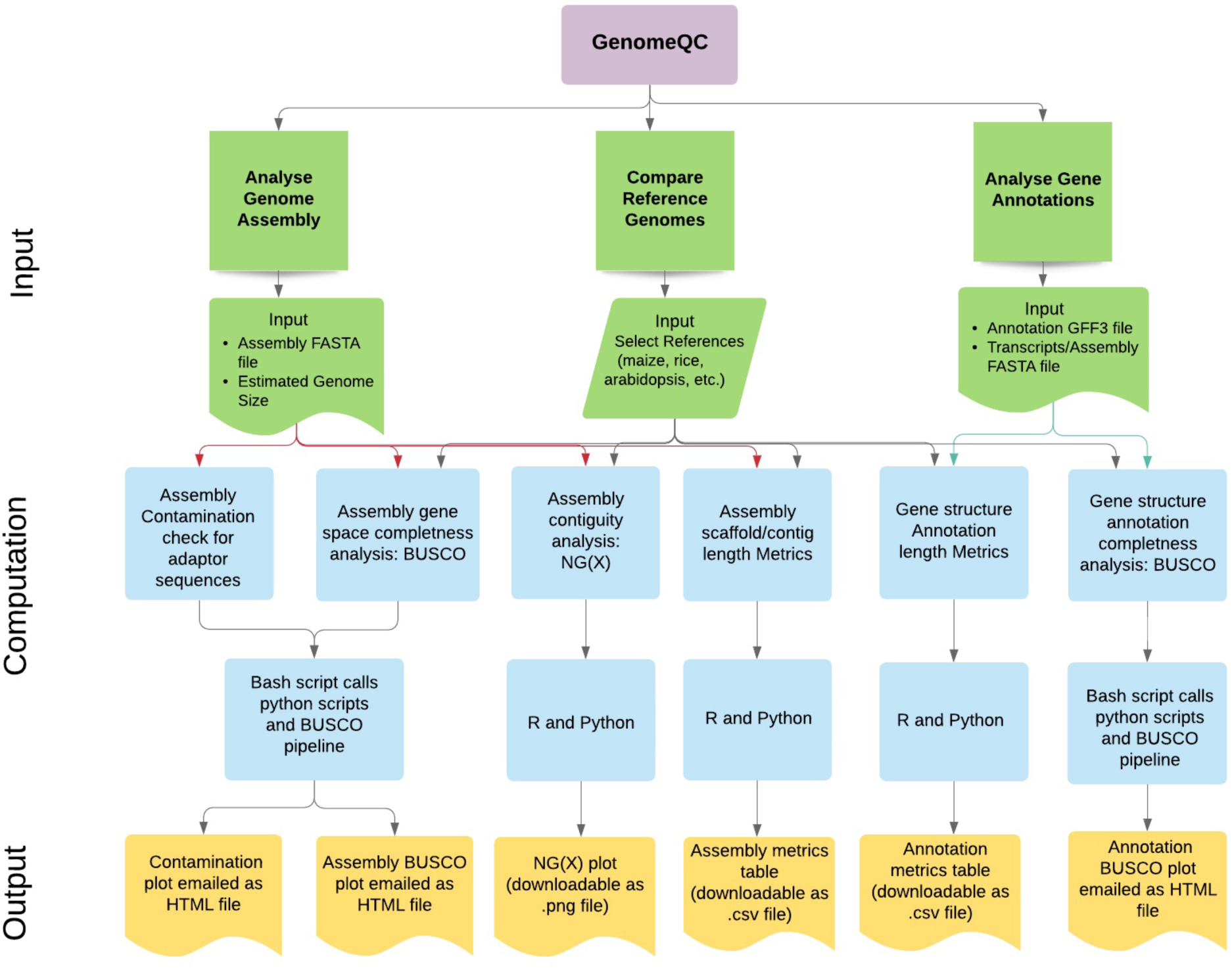
Workflow of the web application. The interface layer of the web application is partitioned into 3 sections: comparing reference genomes, analyzing genome assembly and analyzing gene structure annotations (green). Each of these sections has an input widget panel for files uploads and parameter selection (green). The input parameters and the uploaded data files are then analyzed for contiguity, gene space and repeat space completeness, and contamination check (blue) using bash, R and python scripts (blue) and the different metrics and plots are displayed through the output tabs (yellow).

**Figure 2.**
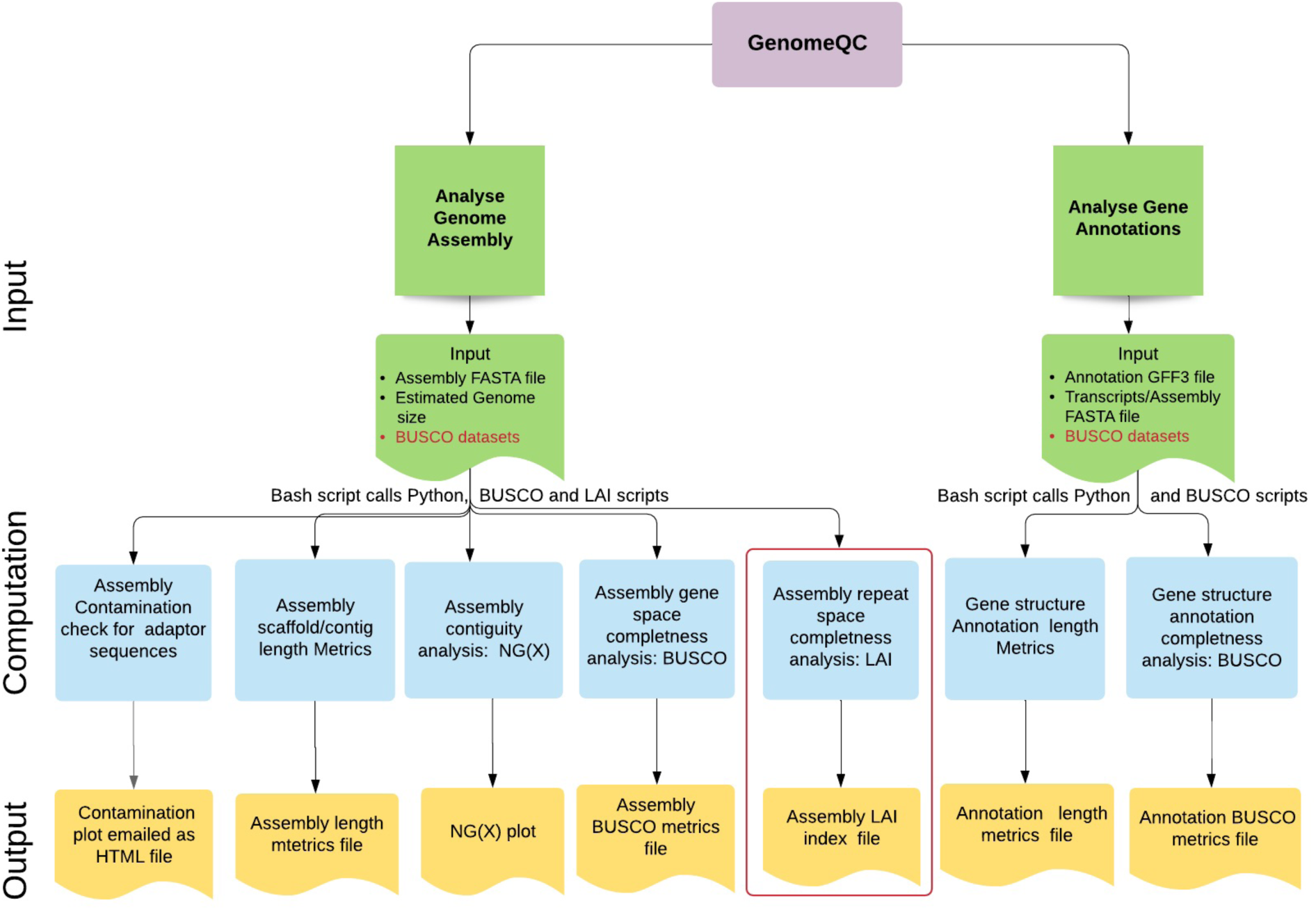
Workflow of the docker image of the GenomeQC pipeline. The containerized version of the GenomeQC pipeline requires BUSCO datasets (highlighted in red) as input in addition to the other input parameters and files (green) required by the web application. Additionally, it allows computation of the LAI index for the input genome assembly (highlighted in red box).

The growing number of assemblies and gene annotations has necessitated the development of metrics that can be used to compare their quality. Such metrics also allow evaluation of the performance of various assembly and annotation methods using the same data. Length metrics (N50/NG50 and L50/LG50 values) provide a standard measure of assembly contiguity [6]. The most commonly reported N50/NG50 values are calculated for the 50% threshold, but NG(X) plots across all thresholds (1-100%) provide a more complete picture [6]. Annotation quality metrics include number of gene models, exons per gene model, and the average lengths of genes, exons and transcripts [7]. Such length and count metrics are useful, but they do not fully capture the completeness of assemblies.

Completeness is better gauged using a set of genes that are universally distributed as orthologs across particular clades of species [8]. A summary of complete single-copy, duplicated, fragmented, and missing Benchmarking Universal Single-Copy Orthologs (BUSCO) genes is often provided as a quantitative measure of genome completeness based on expected gene content. While BUSCO is limited to assessment of the gene space, the LTR Assembly Index [LAI; 9] is capable of gauging completeness in more repetitive genomic regions by estimating the percentage of intact LTR retroelements. LAI is particularly useful for assessing plant genome assemblies, which are often largely comprised of repeats. Recently, dramatic increases in the completeness of repetitive portions of plant genomes have been achieved due to improvements in long-read data [9].

Here, we describe an easy-to-use and interactive web framework based on the R/Shiny package [10] that integrates a suite of quantitative measures to characterize genome assemblies and annotations. Our application, named GenomeQC, provides researchers with a summary of these statistics and allows for benchmarking against gold standard reference assemblies. We have also developed a Docker container of the GenomeQC pipeline that calculates these metrics and supports analysis of large (>2.5Gb) genomes.

### Comparison with similar software programs

Although several tools exist for evaluating and visualizing the quality of genome assemblies like QUAST-LG [11], Icarus [12], LASER [13], REAPR [14], they are challenging to install and configure, do not support assessment of gene structure annotations, and do not determine the completeness of the repetitive fraction of the genome based on LTR retrotransposon content.

GenomeQC provides a user-friendly web framework for calculating contiguity and completeness metrics for genome assemblies and annotations. This tool is unique in that it integrates multiple pipelines so that researchers can obtain a comprehensive assessment of genome and gene model quality. The web application is optimized to compute metrics for small to medium-sized genomes with an upper limit of 2.5 Gb (the approximate size of the maize genome).

GenomeQC also allows researchers to benchmark the analysis relative to gold standard reference genomes. The data input widgets on the side panel of the application include pop-up information icons that provide users with more information on input parameters needed for each analysis. NG(X) and other length metrics can be computed for genomes of any species and require two inputs from the user: a genome assembly or annotation file and estimated genome size. BUSCO (version 3.0.2) analysis requires selection of two additional parameters: BUSCO datasets and AUGUSTUS (version 3.2.1) species [15]. While the web application includes BUSCO and AUGUSTUS options spanning a broad range of species including plants, mammals, bacteria, protists, metazoa, and fungi, comparisons to existing reference genomes are currently tailored to plants.

Additionally, GenomeQC allows a contamination check against the NCBI UniVec database [16] to identify vector and adapter sequences so that these can be removed or masked prior to submission to NCBI or other genome sequence archives.

The containerized version of GenomeQC is configured to additionally compute the LAI value of the input genome assembly. While LAI is a very useful gauge of completeness of the repetitive portion of the genome, it is a computational expressive tool, therefore, only available in the container version of GenomeQC.

### Findings

#### Design concept

GenomeQC is designed to allow users with minimal programming capability to quickly analyze any sequenced genome and to compare assemblies with available reference genomes. Figure 1 and Figure 2 shows the workflow of the web application and the containerized version of the GenomeQC pipeline respectively.

GenomeQC requires two input files and specification of a small number of input parameters. Output is generated as tabular text and stored as comma separated values (CSV). Images are stored as portable network graphic (PNG) files. Output files can easily be downloaded and viewed using Microsoft Excel and other text and image editors.

#### Input files

Two files are required as input for GenomeQC analysis.

“Genome Assembly File” is a sequence file in the standard FASTA format. The file should be gunzipped compressed (.gz) before uploading it to the web-application. The maximum upload limit for the assembly file is 1Gb.

“Genome Structure Annotation File” is a tab separated text file in GFF/GTF format [17]. The file should be gunzipped compressed (.gz) before uploading it to the web-application.

Optional file:

“Transcript FASTA file”: BUSCO analysis of structural annotations requires a transcript file in FASTA format as input. Thus, the user could either directly upload a transcript (DNA nucleotide sequences) file in compressed (.gz) FASTA format or the tool could extract the transcript sequences from the uploaded assembly and annotation files using the gffread utility v0.9.12 [18]. Currently the tool is configured to first use the information from a transcripts file if provided by the user. If the user does not upload the transcripts file, the tool will check whether the sequence IDs in the first column of the GFF file correspond to the headers in the FASTA file. If there is a discrepancy, the tool will print an error message. Otherwise, the BUSCO job will be submitted.

#### Interface Design

The tool’s analysis interface is organized into three sections for three types of analysis.

The “Compare reference genomes” section outputs various pre-computed assembly and annotation metrics from a user-selected list of reference genomes.

The “Analyze your genome assembly” section provides the user the option to perform analysis on their genome assembly as well as benchmark the quality of their genome assembly using precomputed metrics from gold standard reference genomes.

The “Analyze your genome annotation” section provides the user the option to perform analysis on their genome annotations as well as benchmark their analysis versus pre-computed reference genomes.

#### Output Tabs

The **“Assembly NG(X) Plot”** tab calculates NG values for an uploaded assembly based on the input estimated genome size at different integer thresholds (1-100%) and generates a plot showing the thresholds on the x-axis and the corresponding log-scaled scaffold or contig lengths on the y-axis. Genome assemblies with larger scaffold/contig lengths across NG(X) thresholds are more contiguous.

This plot can be downloaded as an image file. The **“Assembly Metrics Table”** and the **“Annotation Metrics Table”** tabs calculate various length and count metrics for the uploaded assembly and annotation files and outputs interactive tables with pop-up plots based on row selection. These tabs provide the user with quick summaries of standard assembly and annotation metrics. These tables can be downloaded as comma separated files.

The **“Assembly BUSCO and Contamination Plots”** tab: calculates and emails BUSCO scores for the uploaded genome assembly and compares it with the pre-computed values of the user-selected reference genomes. A high quality genome assembly is expected to contain a higher number of complete and single copy BUSCO genes (C&S) and a lower number of missing (M) or fragmented (F) BUSCO genes [8]. For contamination analysis, the megablast module of NCBI BLAST+ v2.28.0 [19] is used to identify segments of the assembled genome sequences which may be of vector or adapter origin or from linkers and primer sequences used in cloning cDNA or genomic DNA. Contaminant sequences are downloaded from the NCBI UniVec Database [16]. These plots are emailed as html files which can be opened in a chart studio and customized. The **“Annotation BUSCO plot”** tab calculates and emails the BUSCO scores for the uploaded genome annotations and compares it with precomputed values of the user-selected reference genomes. BUSCO and contamination plots are also emailed as html files. Figure 3 shows the summaries and graphical outputs generated by GenomeQC web application.

**Fig. 3.**
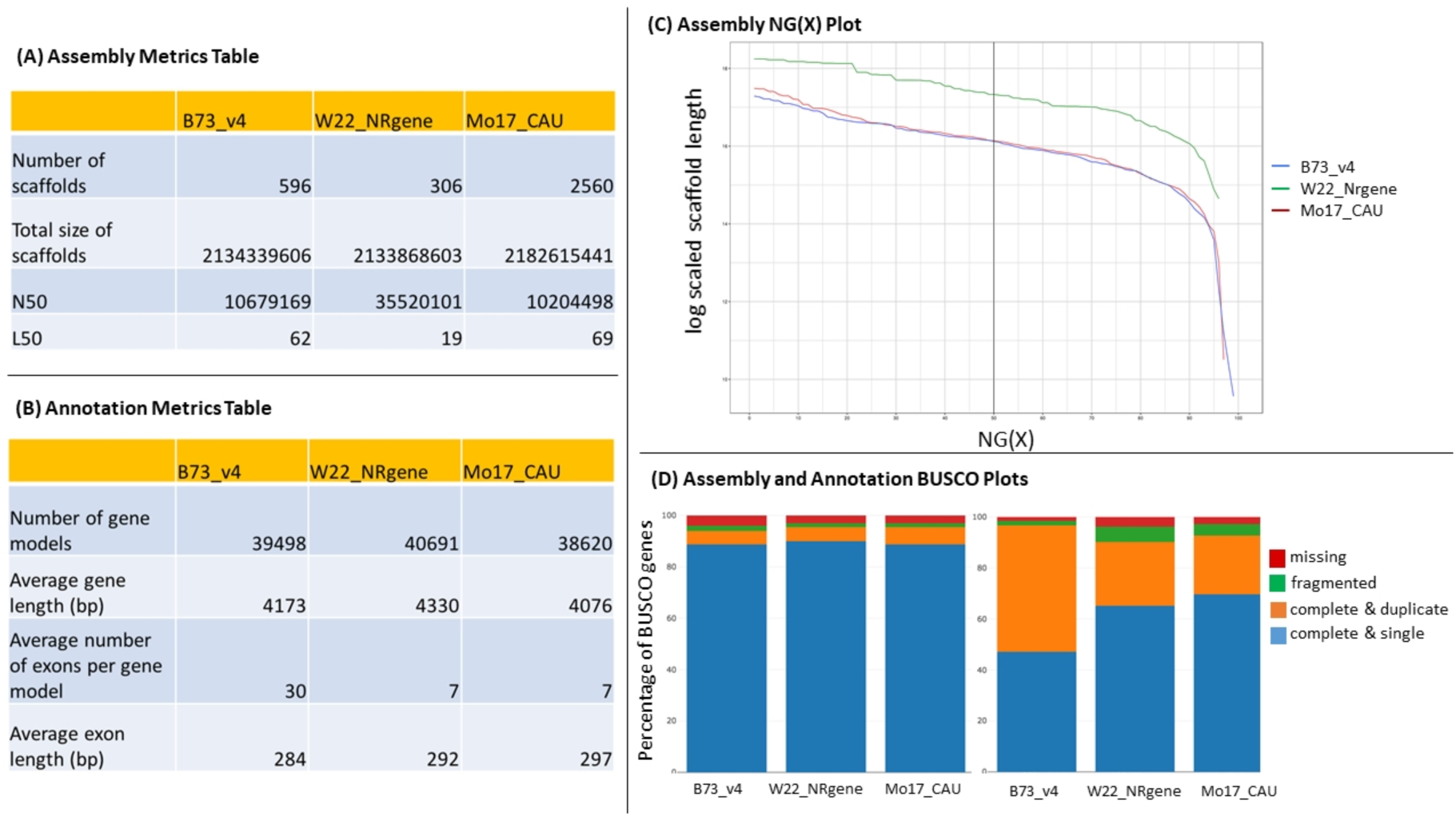
Summaries and graphical output by GenomeQC. Panels (A) and (B) include standard assembly and annotation length metrics generated for maize reference lines B73, W22, and Mo17. Panel (C) is an NG(X) graph in which the x-axis charts NG(X) threshold values (1 to 100%) and the y-axis shows log-transformed scaffold lengths. Each curved line represents scaffold lengths of assemblies at different NG levels with a vertical line at the commonly used NG50 value. Panel (D) shows the relative proportion of complete and single copy (blue), complete and duplicated (orange), fragmented (green), and missing (red) Benchmark Universal Single Copy Ortholog (BUSCO) genes identified for the assembly (left) and gene annotation set (right) of the above mentioned maize lines.

## Discussion

GenomeQC provides a user-friendly and interactive platform for computation and comparison of genome assembly and annotation metrics. The tool has been used to analyze several plant genome assemblies including maize.

Currently, the web application is optimized for analysis of genomes up to 2.5 Gb in size. However, the containerized version of the pipeline available through our GitHub repository can be used to calculate metrics for larger genomes.

## Availability of source code and requirements

Project name: GenomeQC

Project home page:

https://github.com/HuffordLab/GenomeQC

Operating system(s): platform independent Programming language: R, R shiny, Python, Shell script

Other requirements: Docker engine

License: Any restrictions to use by non-academics: None

## Declarations

### List of abbreviations

LTR: Long Terminal Repeats
LAI: LTR Assembly Index
BUSCO: Benchmark Universal Single Copy Orthologs

## Author contributions

NM and CJLD conceived the project. All authors tested the tool, read the manuscript and provided feedback. NM developed the front- and back-end code and was the lead writer for the manuscript. MH, CA, and CJLD were responsible for funding acquisition. MH provided project administration. MRW and AS offered design suggestions and feedback along with running test datasets through GenomeQC. JLP provided network and system administration support.

## Acknowledgements

The authors would like to thank Levi Baber (Iowa State University Director of Research IT) for technical help and Jack Gardiner (Curator at MaizeGDB) for testing the web application and providing helpful suggestions.

