## Supplementary figures and images for "GenomeQC: A quality assessment tool for genome assemblies and gene structure annotations"

### Supplemental Data 1

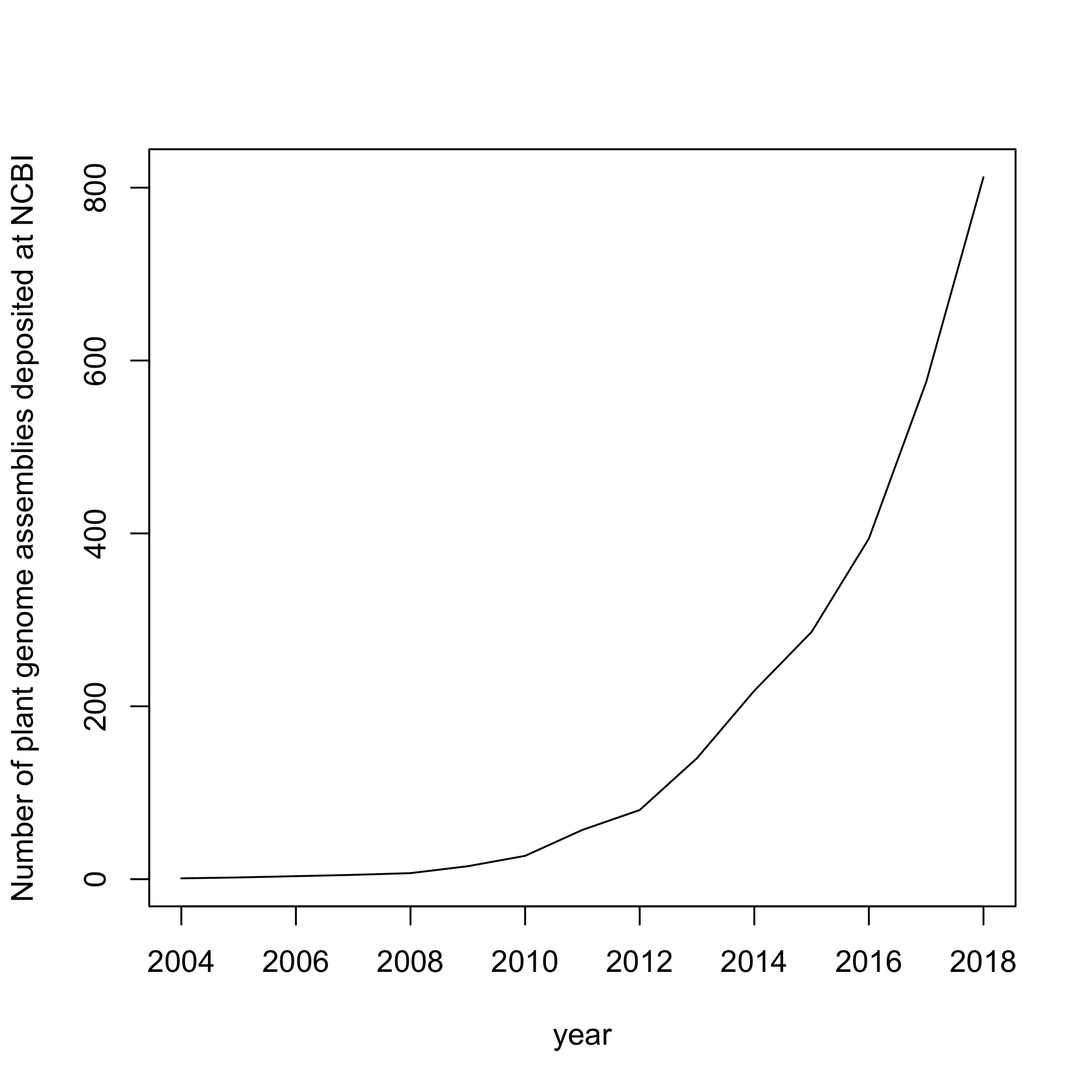

### Supplementary File 1

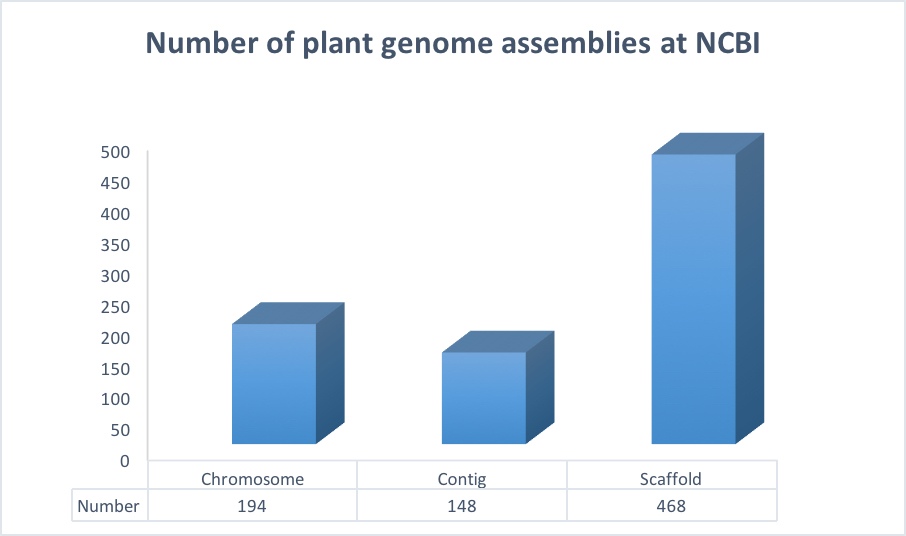
